# hAMRoaster: a tool for comparing performance of AMR gene detection software

**DOI:** 10.1101/2022.01.13.476279

**Authors:** Emily F. Wissel, Brooke M. Talbot, Noriko A. B. Toyosato, Robert A Petit, Vicki Hertzberg, Anne Dunlop, Timothy D. Read

**Affiliations:** Nell Hodgson Woodruff School of Nursing, Emory University, Atlanta, GA, US; Population Biology, Ecology, and Evolution Program, Graduate Division of Biological and Biomedical Science, Emory University, Atlanta, GA, US; Department of Molecular & Biomedical Biology, University of Nebraska at Omaha, Omaha, NE, US; Division of Infectious Diseases, Department of Medicine, School of Medicine, Emory University, Atlanta, GA, USA; Department of Gynecology & Obstetrics, Emory University School of Medicine; Department of Human Genetics, School of Medicine, Emory University, Atlanta, GA, US

**Keywords:** antimicrobial resistance, bioinformatics, metagenomics

## Abstract

**Background:** The use of shotgun metagenomics for AMR detection is appealing because data can be generated from clinical samples with minimal processing. Detecting antimicrobial resistance (AMR) in clinical genomic data is an important epidemiological task, yet a complex bioinformatic process. Many software tools exist to detect AMR genes, but they have mostly been tested in their detection of genotypic resistance in individual bacterial strains. Further, these tools use different databases, or even different versions of the same databases. Understanding the comparative performance of these bioinformatics tools for AMR gene detection in shotgun metagenomic data is important because this data type is increasingly used in public health and clinical settings.

**Methods:** We developed a software pipeline, hAMRoaster (Harmonized AMR Output compAriSon Tool ER; https://github.com/ewissel/hAMRoaster), for assessing accuracy of prediction of antibiotic resistance phenotypes. For evaluation purposes, we simulated a highly resistant mock community and several low resistance metagenomic short read (Illumina) samples based on sequenced strains with known phenotypes. We benchmarked nine open source bioinformatics tools for detecting AMR genes that 1) were conda or Docker installable, 2) had been actively maintained, 3) had an open source license, and 4) took FASTA or FASTQ files as input. hAMRoaster calculated sensitivity, specificity, precision, and accuracy for each tool, comparing detected AMR genes to susceptibility testing.

**Conclusion:** Overall, all tools were precise and accurate at all genome coverage levels tested (5x, 50x, 100x sequenced bases / genome length) in the highly resistant mock community with more variability in the low resistance community (1x coverage). This study demonstrated that different bioinformatic tools and pipelines yield differences in AMR gene identification across drug classes, and that these differences become important if researchers are interested in resistance to specific drug classes.

**Significance:** Software selection for metagenomic AMR prediction should be driven by the context of the clinical/research questions and tolerance for true and false negative results. The ability to assess which bioinformatics tool best fits a particular dataset prior to beginning a large-scale project allows for more efficient processing and analysis using optimal tools for a particular research question. As prediction software and databases are in a state of constant refinement, the approach used here—creating synthetic communities containing taxa and phenotypes of interest along with using hAMRoaster to assess performance of candidate software—offers a template to aid researchers in selecting the most appropriate strategy at the time of analysis.

## Introduction

Antibiotic resistant infections pose a serious threat not only to public health but to the agricultural, veterinary, and food safety industries. The misuse of antibiotics in healthcare and livestock production has led to widespread antimicrobial resistance in diverse environments and has emerged as a threat to global health.^1 2^ The burden of multi-drug resistant pathogens is increasing globally, creating complex clinical scenarios in which there are limited (if any) therapeutic options, resulting in increased mortality and healthcare costs for common medical procedures.^3^ Genes that confer antimicrobial resistance (AMR) are increasingly present in commensal members of the human microbiome and are recognized as an important reservoir for conferring pathogen resistance through horizontal gene transfer.^4,5^

Two key approaches to mitigating AMR infections are antibiotic stewardship and AMR surveillance. While antibiotic stewardship focuses on using antibiotics appropriately, AMR surveillance focuses on describing AMR genes already present in a community. Currently, AMR surveillance typically relies on phenotypic characterization through culture or genotypic characterization through molecular diagnostics based on PCR and hybridization techniques.^6^ However, there is a move toward genome-based methods ^7^ with the Illumina short-read platform being the dominant platform for data generation at the present time.^8^

Sequencing technology has revolutionized research across many disciplines, with more applications found every year as both the technologies and analysis methods advance. This is particularly evident in the use of metagenomic data for the microbial surveillance of antimicrobial resistance (AMR), as microbial communities can be characterized without the need to first isolate and culture the specimen prior to analysis.^9–11^ As the cost and time of sequencing has dramatically decreased, petabytes of data are quickly generated, with Illumina short reads becoming more prevalent.^8,12,13^ Detecting AMR genes potential through non-culture based, high throughput DNA sequencing and bioinformatic approaches is of growing relevance and importance.

There are many bioinformatic tools created to process large amounts of data while following open-science principles.^14^ Open science is a term used to describe data that is Findable, Accessible, Interoperable, and Reusable or (FAIR) and that are open-source.^15^ With so many options available, it is important that investigators determine which open-source tool is the best suited for their research question. One way to address issues with replicability and variance across studies is to establish standardized bioinformatics pipelines and best practices, as has been done, for example,by the National Microbiome Data Collaborative (NMDC).^16^ However, for many researchers, a standardized bioinformatics pipeline may not the best suited for their data or research question.^14^

As shotgun metagenomic sequencing is emerging as a powerful tool for detecting AMR,^17^ it is essential to evaluate how well different tools perform. In addition to testing AMR gene prediction tools against widely available metagenome samples, they should be compared in samples with extensive phenotypic resistance (acquired and mutational AMR genes). Here, we describe a software pipeline, hAMRoaster, that provides metrics on tool performance in detecting AMR genes from known resistant phenotypes and can therefore help in decision-making about which tools will be adequate for detecting resistance to the drug classes being studied.

## Methods

For a schematic overview of the methods, see **Figure One.**

### Development of a software pipeline, hAMRoaster, to assess results of antibiotic resistance prediction

hAMRoaster was written as a conda installable command line tool in a Python script and requires three inputs: a) the text output of AMR tool on a FASTQ or FASTA test file, such as a text file processed through hAMRonization,^18^ b) a list of known phenotypes associated with the test file or samples names, and c) (optional) a tab formatted table which matches antibiotic drugs with their drug class. If option c) is not specified a default table is used. The output of the program is a set of performance metrics that include sensitivity and specificity. A conda installable version of the software was deposited in the Bioconda^19^ database. The Github site for the software is https://github.com/ewissel/hAMRoaster.

hAMRoaster requires, as input, a formatted results table of runs by AMR detection tools. This table is identical to that produced by the hAMRonization^18^ software. hAMRonization is conda installable and can compile the output of many AMR tools into a unified format. shortBRED^20^ and fARGene^21^ are not included in hAMRonization at the time of analysis, so hAMRoaster can take the path to the raw output for these tools and partially match it to the hAMRonization output.

hAMRoaster requires an input to the “known” phenotypic resistance in the mock community (--AMR_key flag of hAMRoaster), such as a result of susceptibility testing tables that are available from NCBI Biosamples. Antibiotics in the table of known phenotypic resistances are matched to their respective drug classes. Results classified as “susceptible” in susceptibility testing are considered “susceptible”, and “intermediate” results are ignored. In cases where susceptibility testing occurred with two or more agents, each agent is considered independently (e.g. resistance to “amoxicillin-tetracycline” is treated as resistance to “amoxicillin” and “tetracycline” independently). Each identified AMR gene is labeled with its corresponding drug class for comparison. In instances where a gene confers resistance to multiple drug classes, the detected gene is split into multiple rows so that each conferred resistance can be independently compared to the susceptibility testing. Gene to drug class linkage is verified using the CARD database^22^ when applicable by accession ID. Any genes corresponding to ‘unknown’ or ‘other’ drug classes (including hypothetical resistance genes) are excluded from further analysis. Genes that confer resistance to an antibiotic that is only administered and effective in combination with another drug (e.g. clavulanic acid in amoxicillin-clavulanic acid) are classified as ‘Other’ and excluded from analysis.

A detected AMR gene is labeled as a true positive by hAMRoaster if the drug class matched to an AMR gene corresponds to a drug class that tests “resistant” in the susceptibility testing for the mock community. Similarly, a false positive is coded as a drug class that is called by the software, but tested as susceptible in the mock community (--AMR key parameter). Observed AMR genes are labeled “unknown” if the corresponding drug class is not tested in the mock community and is not included in the AMR key file. Once true/false positives and true/false negatives are determined per tool, hAMRoaster calculates sensitivity, specificity, precision, accuracy, and percent unknown.

### Creation of multiple synthetic mock communities of antibiotic resistance bacteria

#### *Simple synthetic community with* high *resistance*

Bacterial members of the base mock community were chosen from NCBI’s BioSample Database^23^ and met the following criteria: (1) the strain had extensive antibiotic susceptibility testing data using CLSI or EUCAST testing standards as part of the public NCBI BioSample record; (2) the strain was isolated from human tissue; (3) the strain was the cause of a clinical infection; (4) the FASTA was available to download from NCBI BioSample Database.^23^ Eight bacteria, each representing a different species, with overlapping resistance to 43 antibiotics across 18 drug classes, were selected for the mock community (**Table 1**). The included taxa were *Acinetobacter baumannii* MRSN489669, *Citrobacter freundii* MRSN12115, *Enterobacter cloacae* 174, *Escherichia coli* 222, *Klebsiella pneumoniae* CCUG 70742, *Pseudomonas aeruginosa* CCUG 70744, *Neisseria gonorrhoeae* SW0011, and *Staphylococcus aureus* LAC (Table 1).

**Table 1A:** Clinical isolates included in the high resistance simulated community. (susceptibility test is in the spreadsheet, will have to be supplemental bc so big)

| Strain | Testing Standard<br>(CLSI or EUCAST) | BioSample ID | Link |
| --- | --- | --- | --- |
| *Neisseria gonorrhoeae*<br>SW0011 | CLSI | SAMN15960549 | https://www.ncbi.nlm.nih.gov/biosample/SAMN15960549 |
| *Klebsiella pneumoniae*<br>CCUG 70742 | EUCAST | SAMN07602587 | https://www.ncbi.nlm.nih.gov/biosample/SAMN07602587 |
| *Pseudomonas aeruginosa* CCUG<br>70744 | EUCAST | SAMN07602569 | https://www.ncbi.nlm.nih.gov/biosample/SAMN07602569 / |
| *Acinetobacter baumannii*<br>MRSN489669 | CLSI | SAMN12087686 | https://www.ncbi.nlm.nih.gov/biosample/SAMN12087686 |
| *Enterobacter cloacae*<br>174 | CLSI | SAMN04456586 | https://www.ncbi.nlm.nih.gov/biosample/SAMN04456586 |
| *Citrobacter freundii*<br>MRSN12115 | CLSI | SAMN13412315 | https://www.ncbi.nlm.nih.gov/biosample/SAMN13412315 |
| *Staphylococcus aureus*<br>LAC | CLSI | SAMN08391108 | https://www.ncbi.nlm.nih.gov/biosample/SAMN08391108 |
| *Escherichia coli* 222 | CLSI | SAMN05194390 | https://www.ncbi.nlm.nih.gov/biosample/SAMN05194390 |

**Table 1B:**
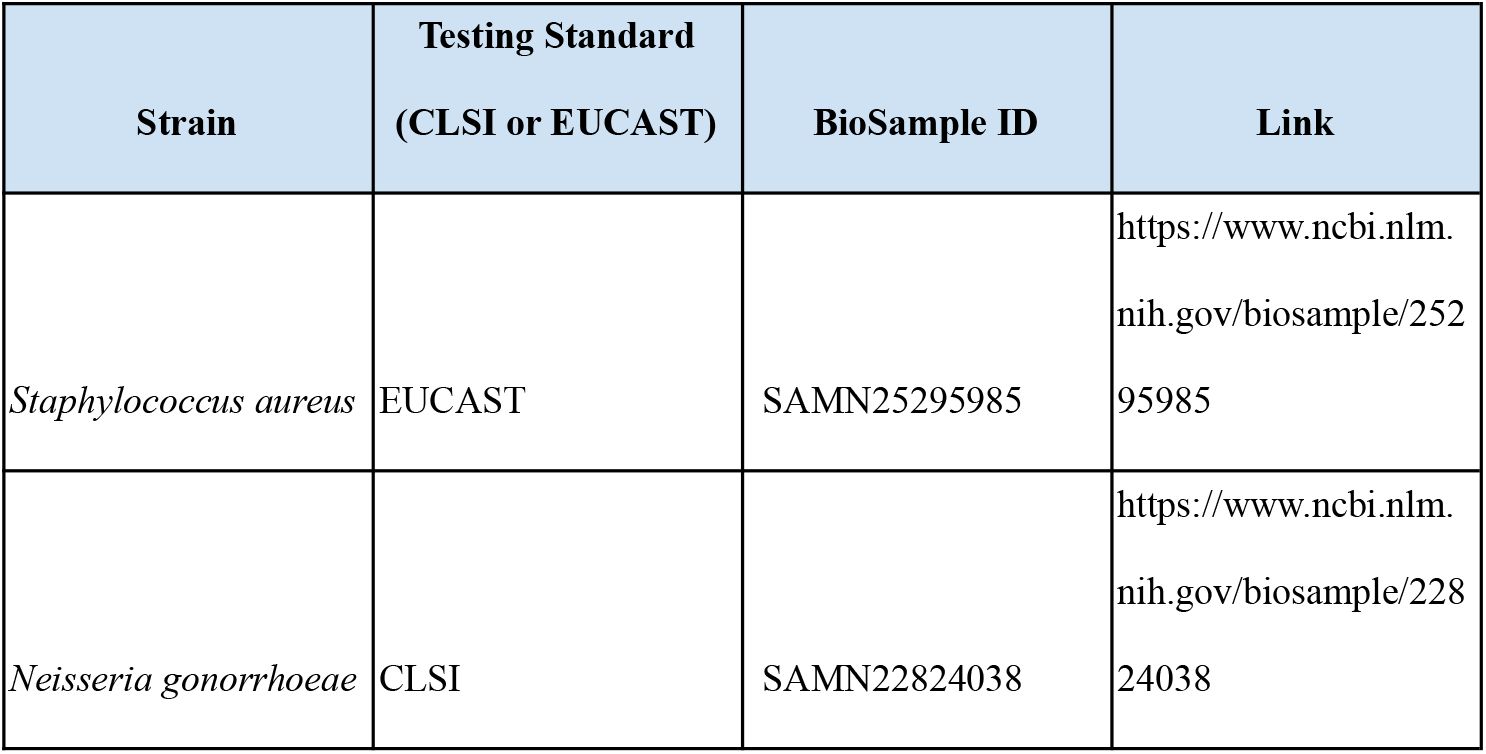
Clinical isolates included in the low resistance simulated community. (susceptibility test is in the spreadsheet, will have to be supplemental bc so big)

Paired-end FASTQs were simulated by NCBI’s ART ^24^ using default parameters for HiSeq 2500 at three levels of average sequence coverage (5x, 50x, and 100x sequenced bases / genome length) and are available on FigShare (https://figshare.com/account/home#/projects/125974). Simulated FASTQs were subsequently concatenated to resemble shotgun metagenomics reads, and metaSPAdes^25^ was used to create assembled contigs. The FASTQs were simulated with approximately equal numbers of reads of each genome.

### Complex synthetic clinical mock community with low resistance

We created a community profile with previously simulated human metagenomes ^26^ and added a single AMR isolate collected from a human infection at 1x coverage to simulate a human metagenome with restrictive phenotypic resistance. We included samples 0 through 5 from CAMISIM,^26^ a set of previously simulated human metagenomes, and combined these with simulated fastqs from one of two isolates from human infections, SRR17789825^27^ for even sample numbers and SRR16683675^28^ for odd sample numbers.

### Running antibiotic prediction software on mock communities

All tools for AMR prediction were run on the mock community and restrictive samples at all coverage levels using default settings for either simulated FASTQ or assembled contigs. Default settings were used as it is what most users use and understand to be the developer recommendations. When both options were available, assembled contigs were run.

### Statistical Analysis

Data were analyzed in Python v3.7.7 and plotted in R v4.0.4. hAMRoaster calculated all performance metrics reported in Table 3. Unweighted Cohen’s kappa was calculated using R package IRR^29^ for each pairwise combination of tools to test agreement between tools.

**Table 2:**
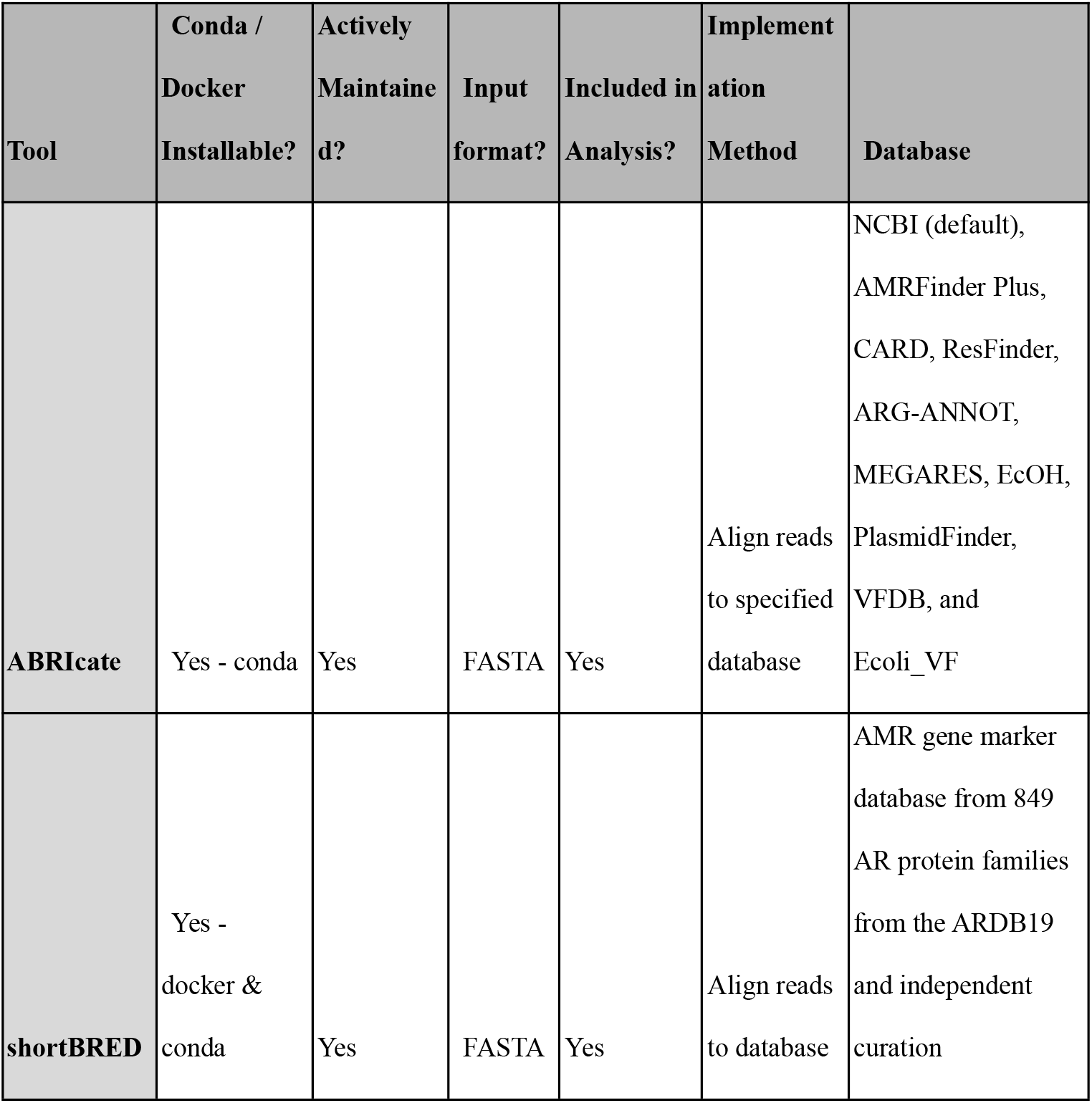
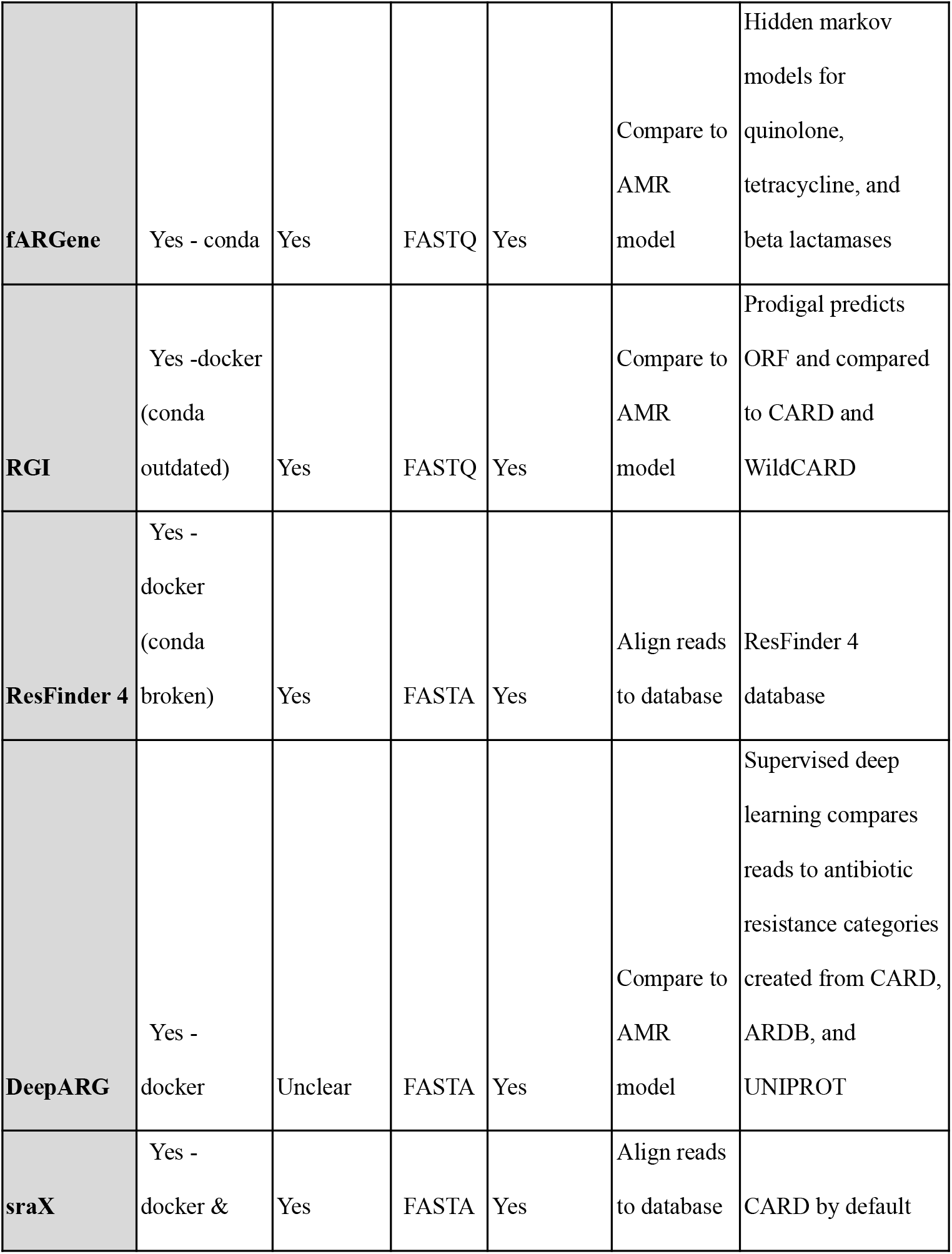
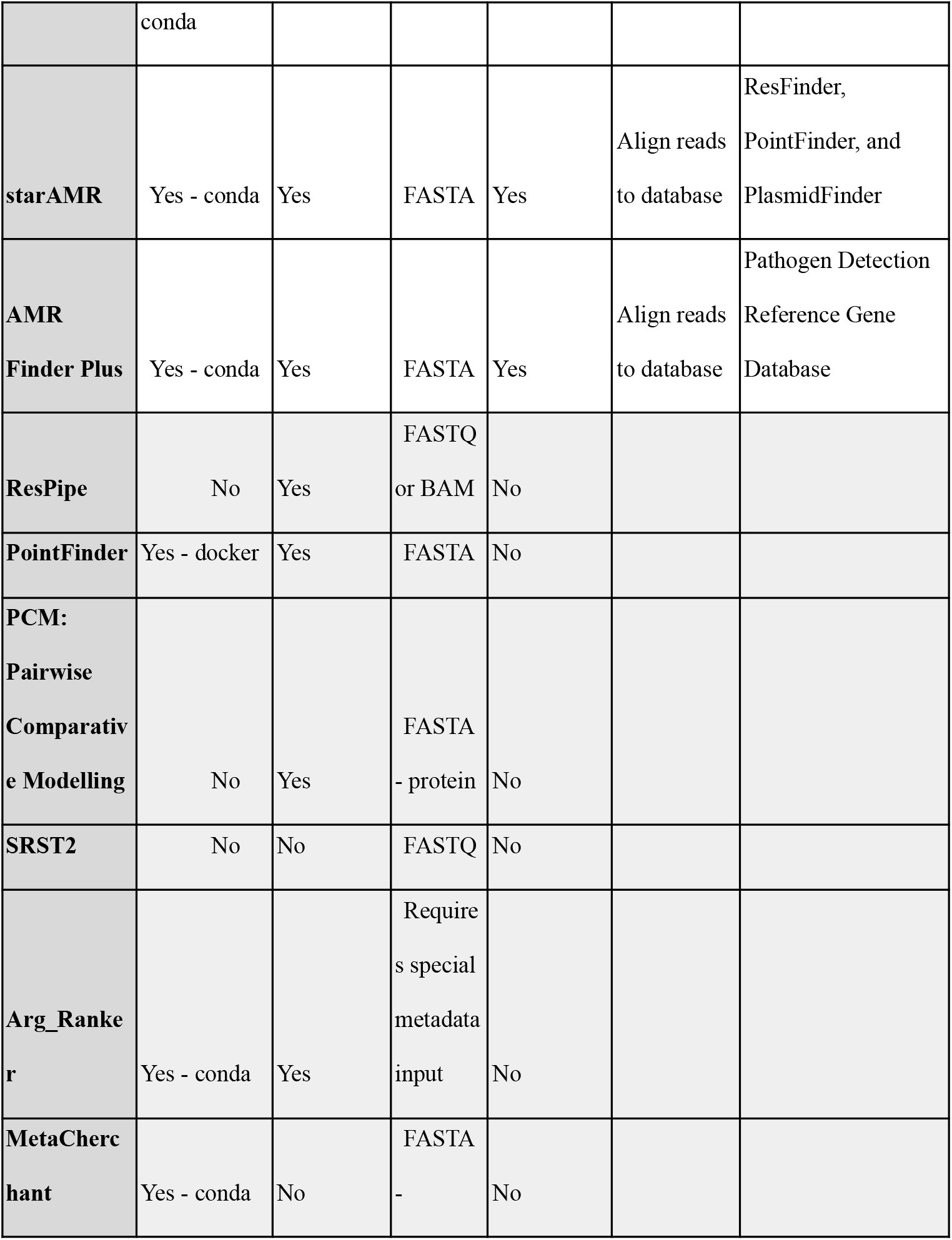
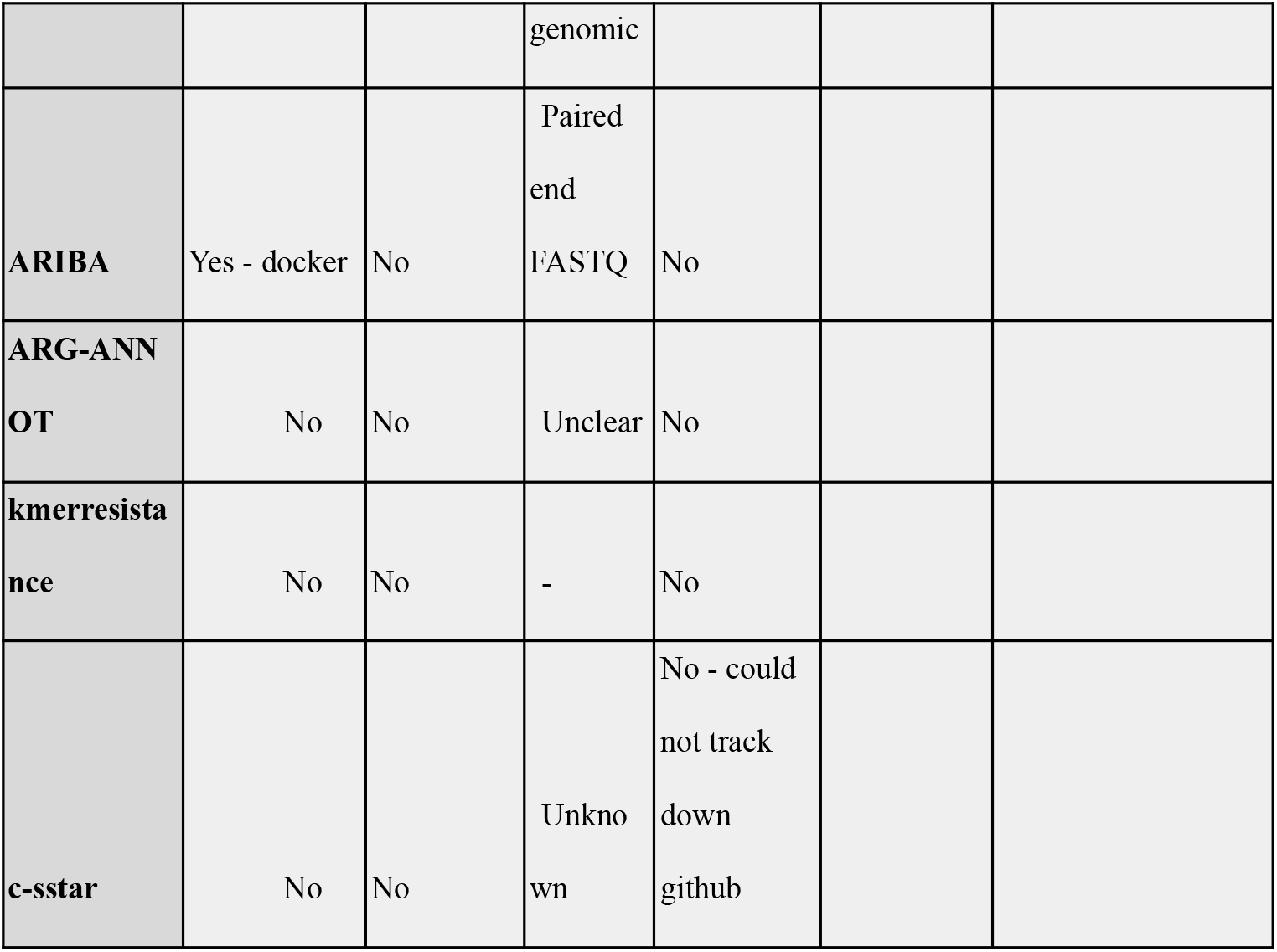
Tools identified from search methods with the selection criteria and whether they subsequently worked or not.

**Table 3A:**
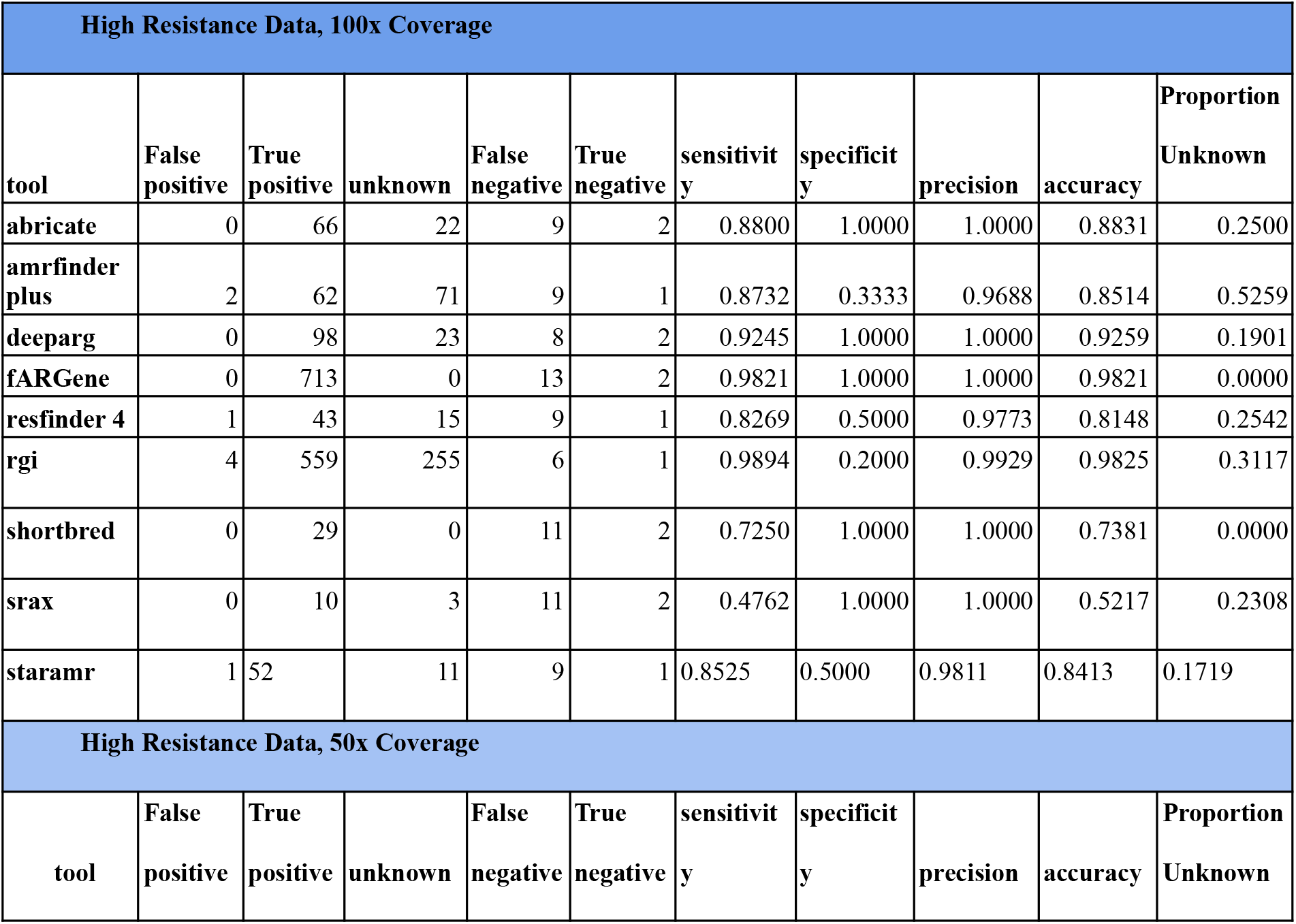
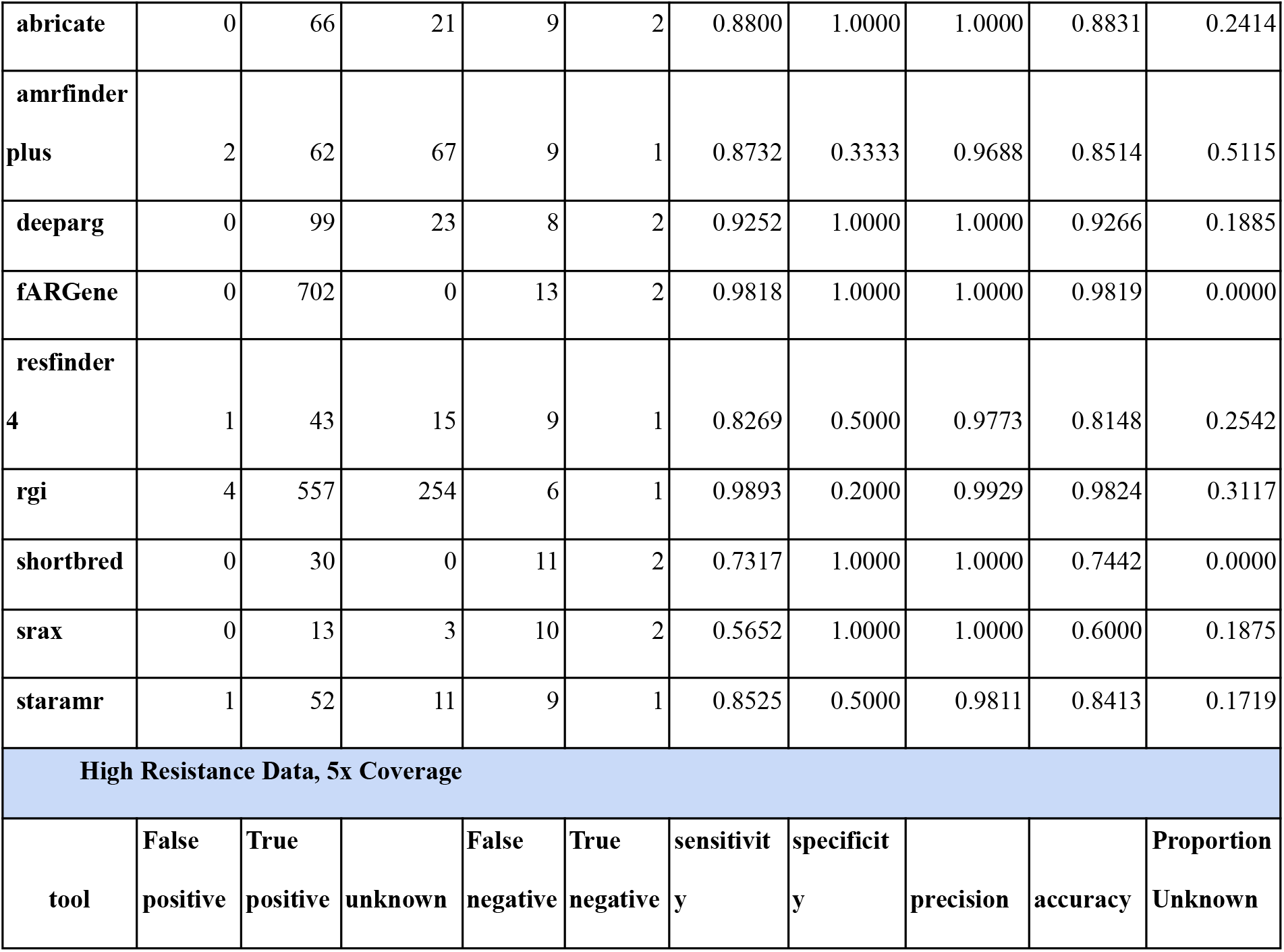
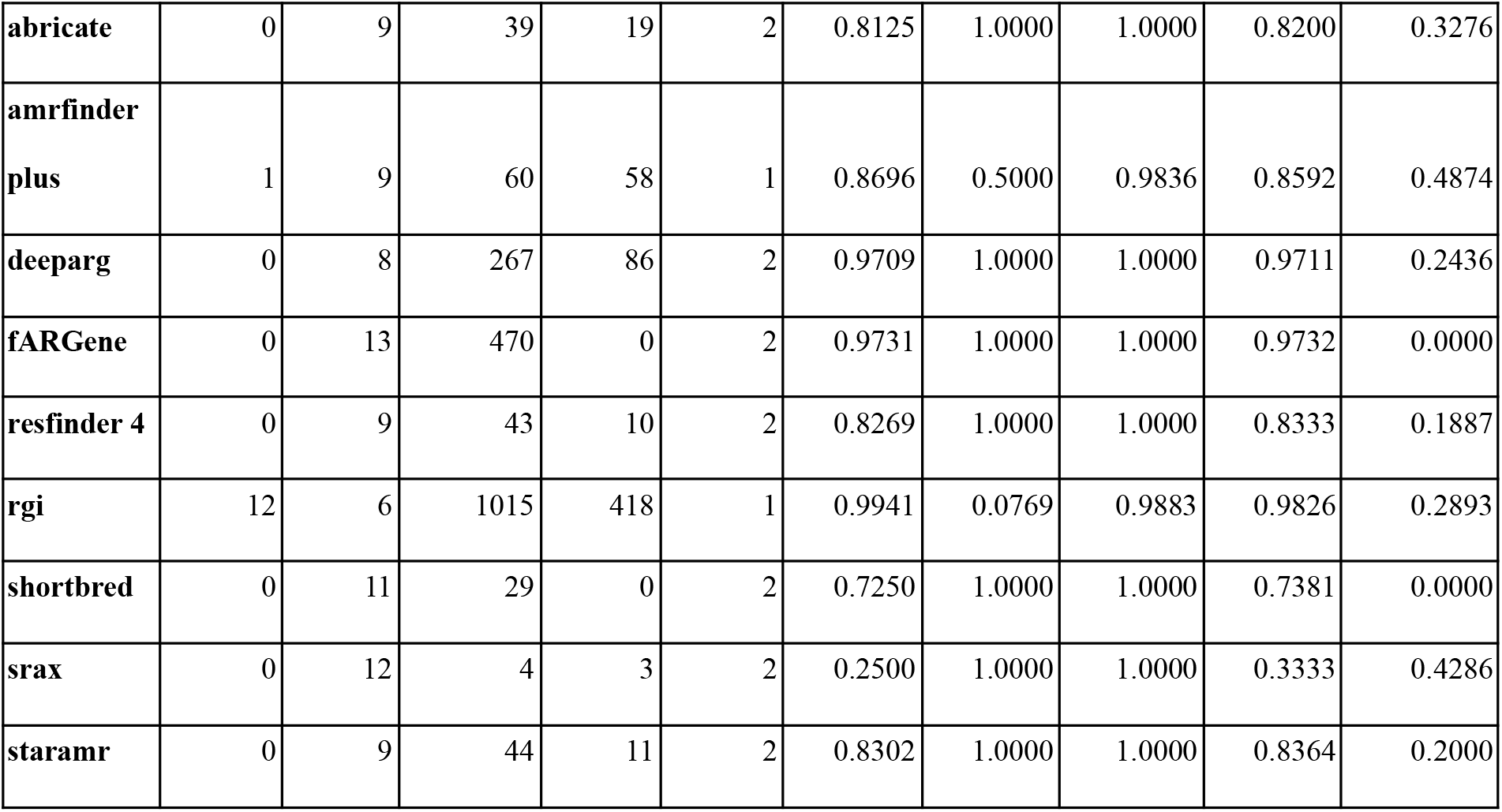
Summary Statistics for the high resistance data from hAMRoaster: These are the counts and metrics as calculated by the hAMRoaster pipeline. Formulas for all metrics are as follows:

Specificity = TN / (TN + FP)
Sensitivity = TP / (TP + FN)
Precision = TP / (TP + FP)
Accuracy = (TP + TN) / (TP + FP + TN + FN)
Proportion Unknown = unknown / (TP + FP + unknowns)

**Table 3B:**
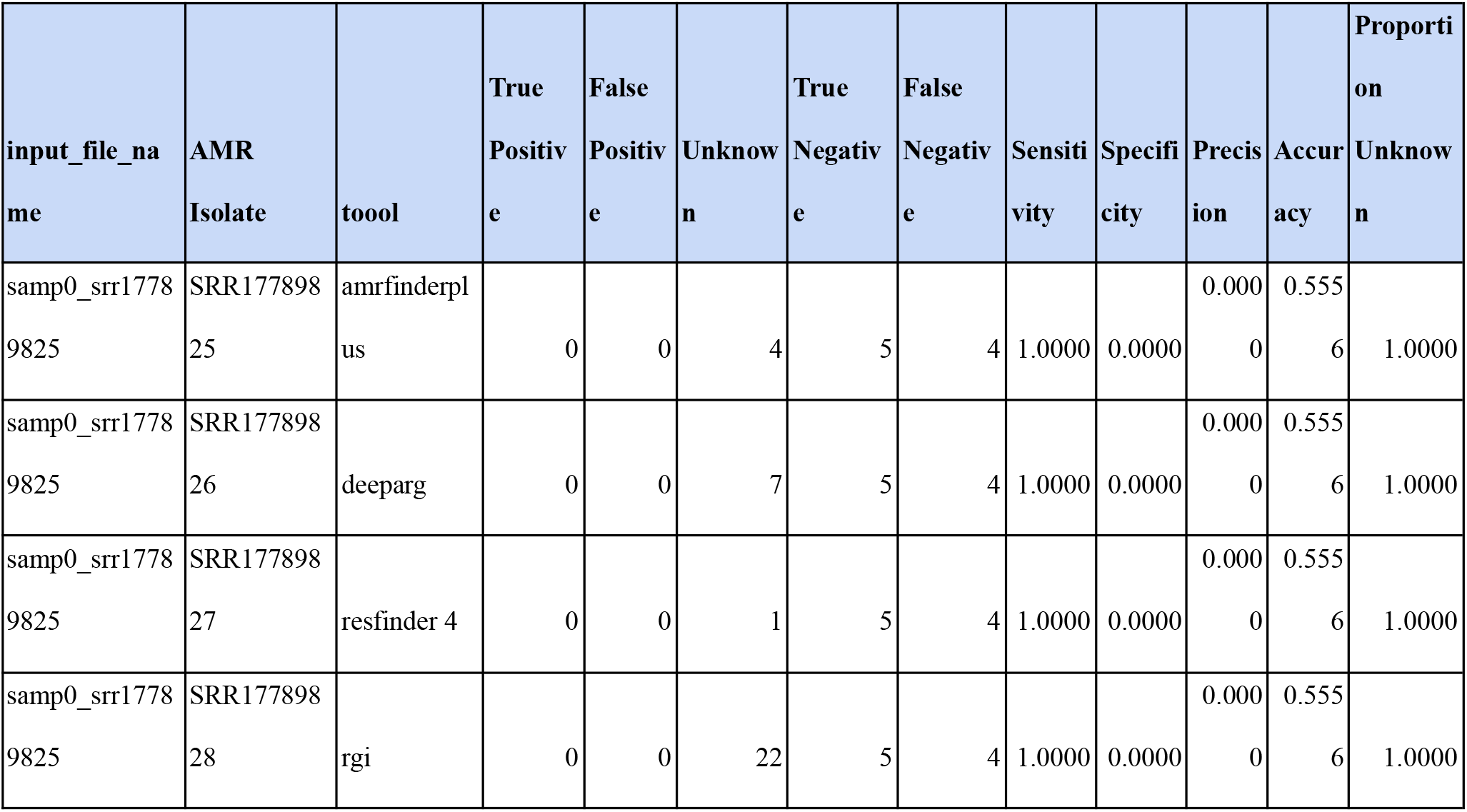
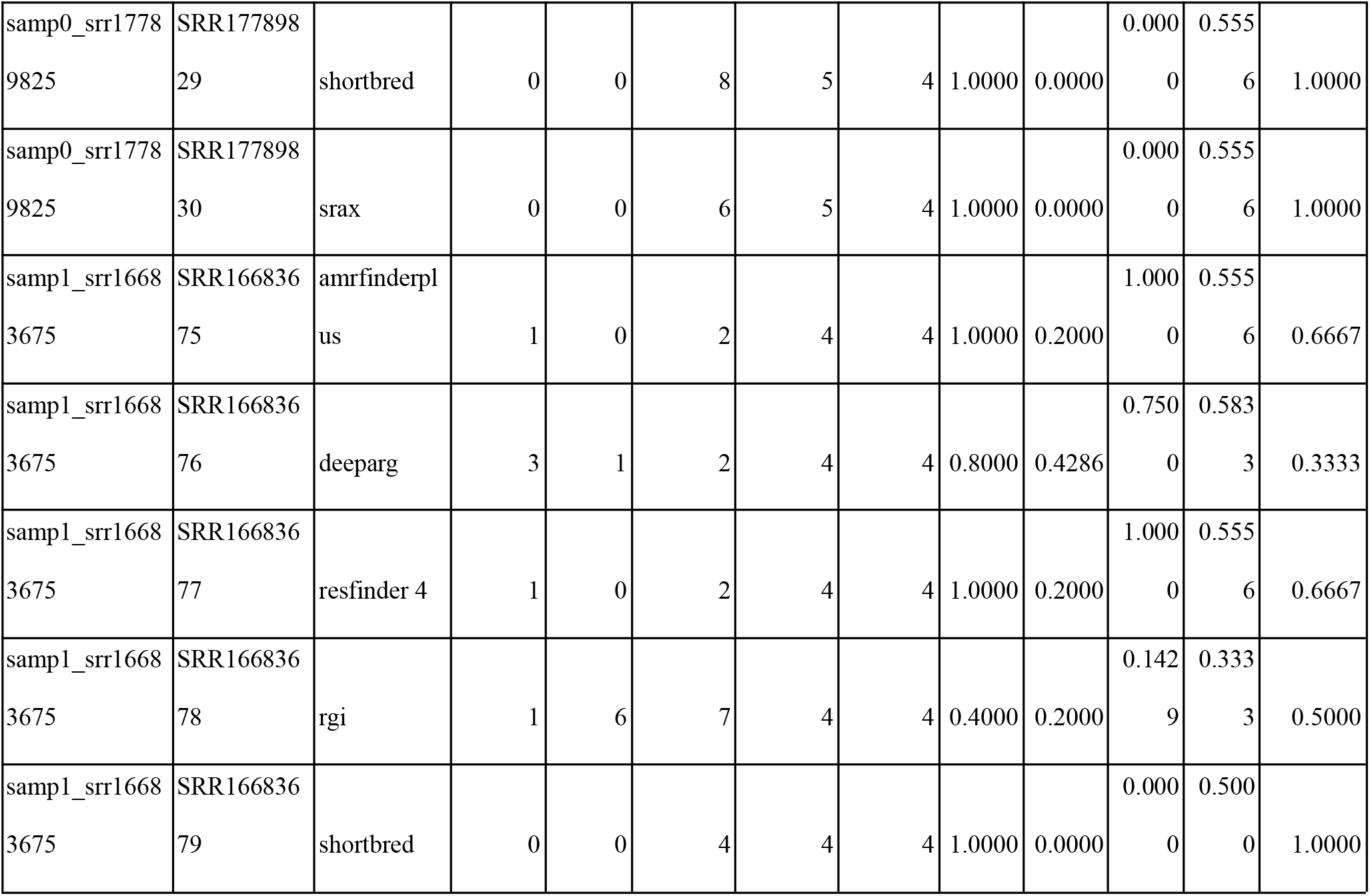
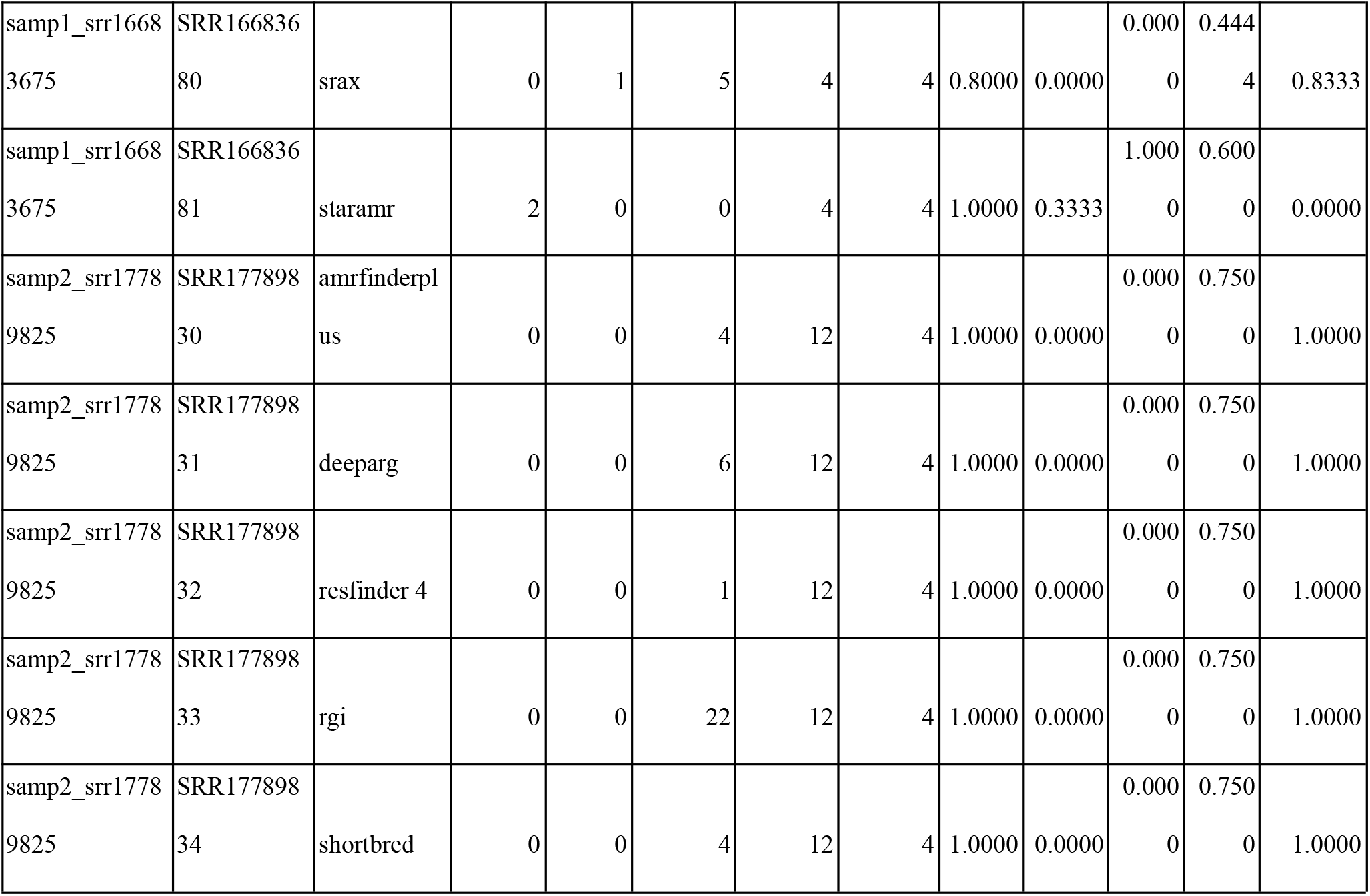
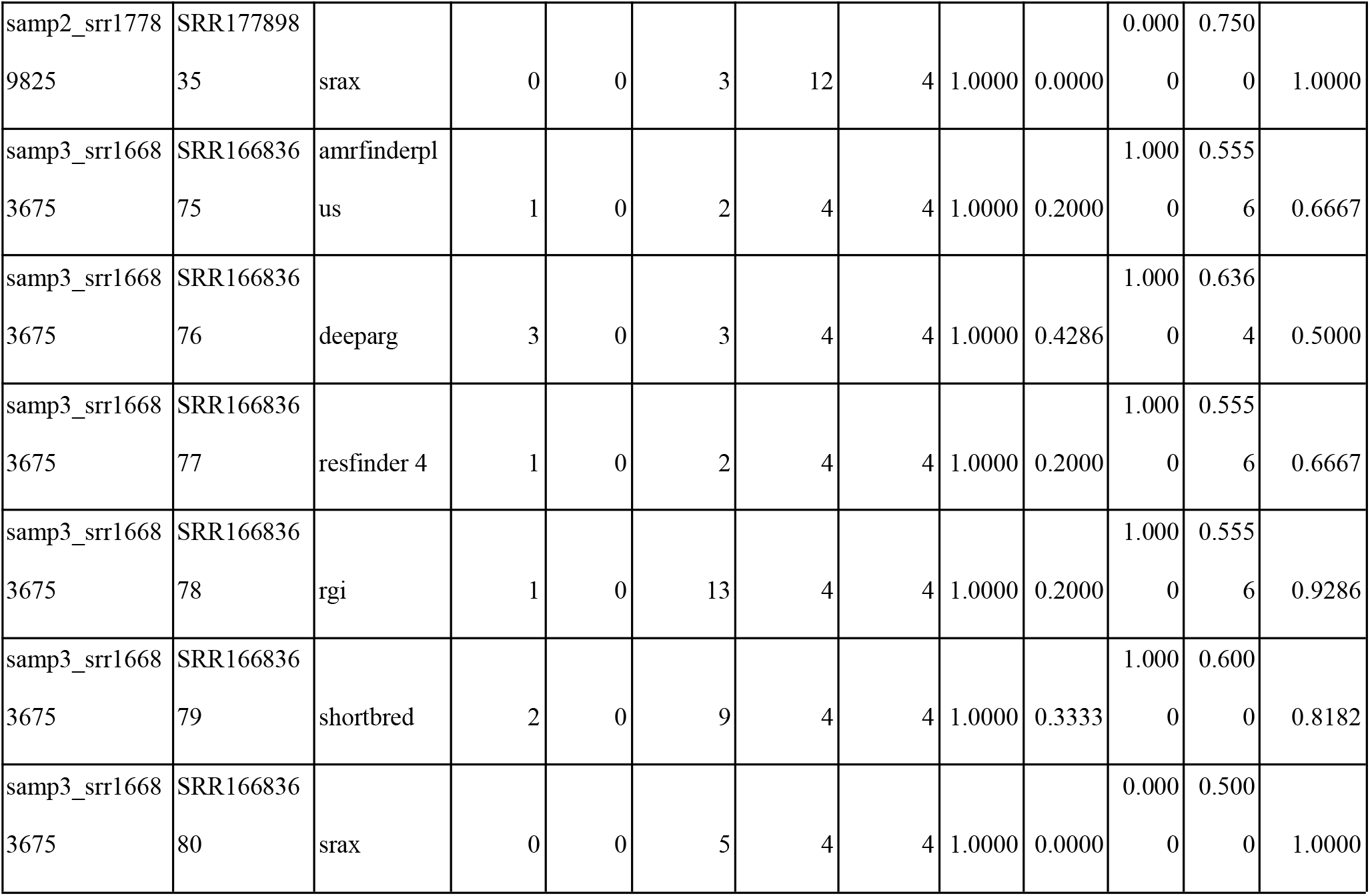
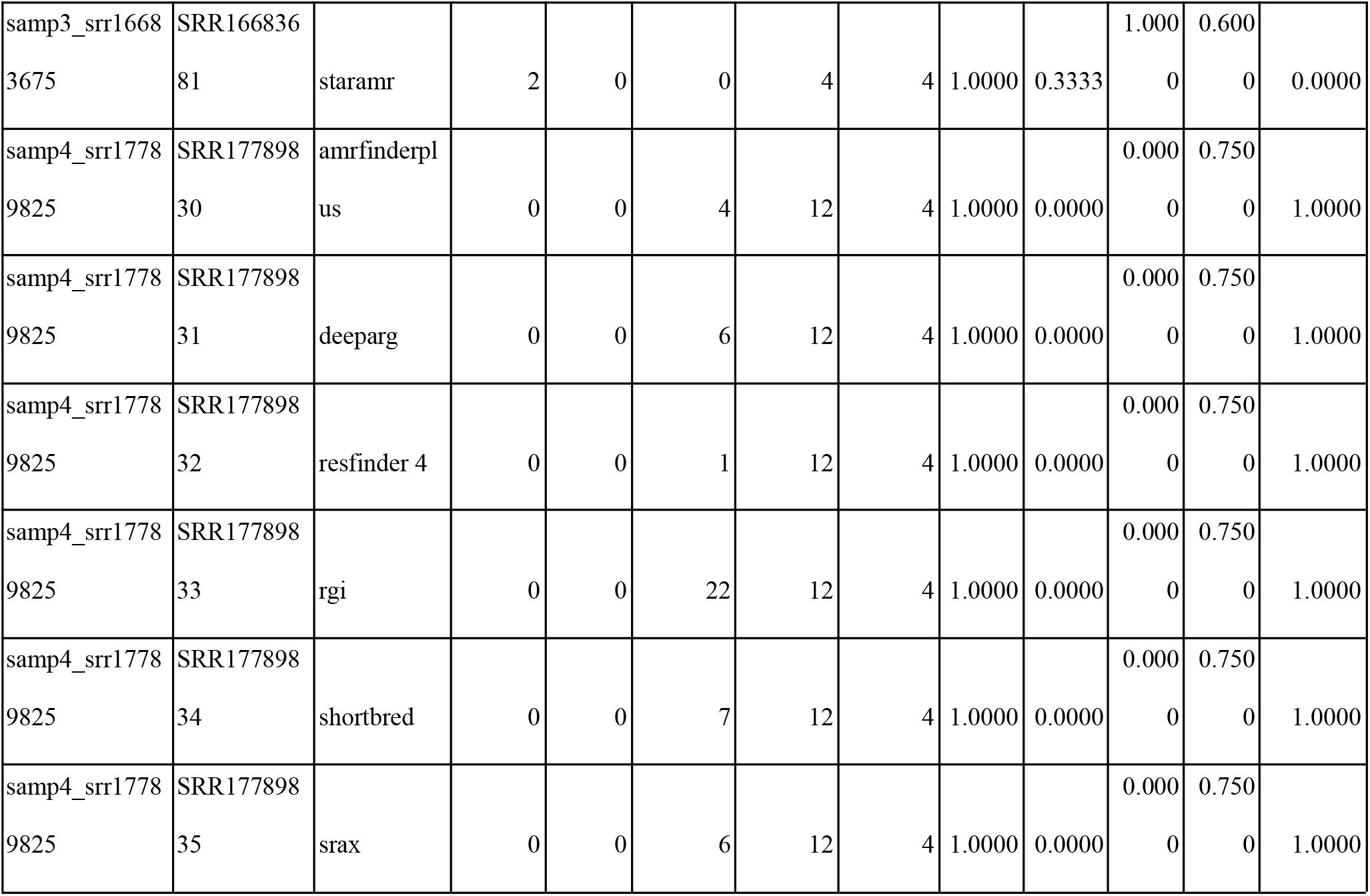
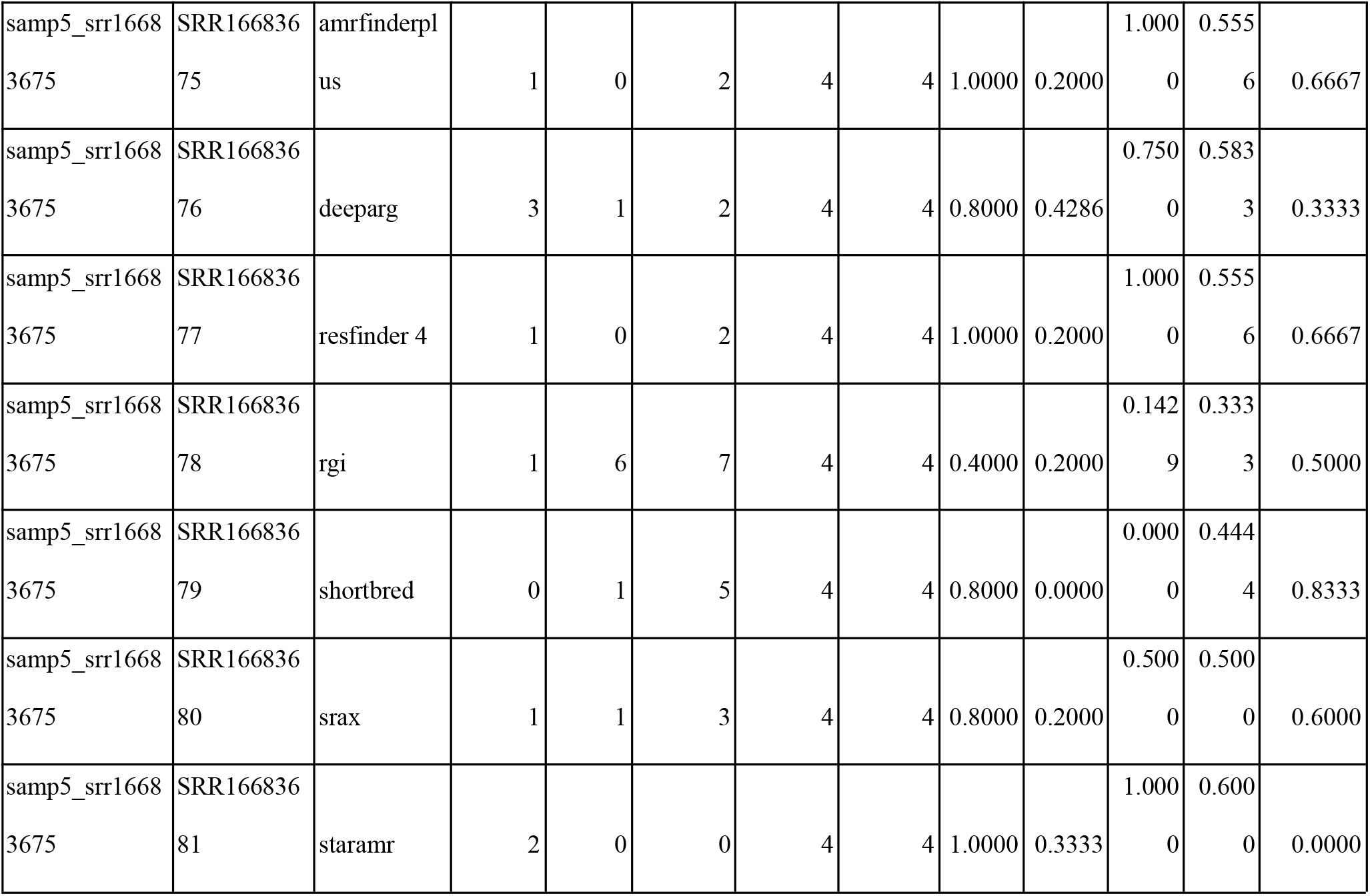
Summary Statistics for the low resistance data from hAMRoaster: These are the counts and metrics as calculated by the hAMRoaster pipeline.

### Data Availability

All data and code is available on the hAMRoaster GitHub repository (https://github.com/ewissel/hAMRoaster) and figshare (for large files; https://figshare.com/account/home#/projects/125974)

## Results

### Selection of nine open source, conda-installable tools for detection of antibiotic resistance phenotypes

To identify tools for antibiotic resistance prediction, we used a multi-headed search strategy. We searched PubMed using terms “AMR”, “antibiotic resistance genes”, “bioinformatics”, and “antimicrobial resistance”. We also searched GitHub using the same set of terms. Once an initial list of tools was compiled, we performed a second PubMed literature review including the search terms from above plus the names of the tools (“tool 1” OR “tool 2”). We also used Twitter to ask the research community what bioinformatic tools they use to identify AMR (supplementary text 1). These searches identified 16 potential tools to identify AMR genes (**Table 2**). The search for tools concluded on March 1, 2021.

For an identified tool to be considered eligible for comparison, it had to meet the following criteria: (1) be conda or Docker installable; (2) have source code publicly available in a data repository and be actively maintained (defined as tool updates or GitHub responses within the last year); (3) have an open source license; and (4) take FASTQs or FASTAs as input files. Nine tools met the criteria to be included in this analysis: ABRIcate ^30^, fARGene ^31^ ResFinder ^32^, shortBRED^20^, RGI ^33^, AMRFinderPlus ^34^, starAMR ^35^, sraX ^36^, and deepARG ^37^. PointFinder also qualified^38^, but was a subtool of ResFinder and only identified mutational resistance for some organisms, so it was excluded from analysis. The code used to install and run all tools is available on the hAMRoaster GitHub.

### ABRIcate

ABRIcate v.1.0.1 took contig FASTA files as inputs and compared reads against NCBI AMRFinder Plus^34^ by default, though there are options to compare against CARD,^33^ ResFinder,^32^ ARG-ANNOT,^39^ MEGARES,^40^ EcOH,^41^ PlasmidFinder,^42^ VFDB,^43^ and Ecoli_VF,^44^ which are also pre-downloaded. ABRIcate reported on acquired AMR genes and not mutational resistance.

### shortBRED

shortBRED^20^ v0.9.3 used a set of marker genes to search metagenomic data for protein families of interest. The bioBakery^45^ team published an AMR gene marker database built from 849 AR protein families derived from the ARDB ^46^ v1.1 and independent curation alongside shortBRED, which is used in this study.

### fARGene

fARGene^21,31^ v.0.1 used Hidden Markov Models to detect AMR genes from short metagenomic data or long read data. This was a different approach from most other tools which compare the reads directly. fARGene has three pre-built models for detecting resistance to quinolone, tetracycline, and beta lactamases, which were tested in this study. fARGene can predict unknown ARGs using its gene models.

### RGI

RGI^33^ v5.1.1 used protein homology and SNP models to predict ‘resistomes’. It used CARD’s protein homolog models as a database. RGI predicts open reading frames (ORFs) using Prodigal,^47^ detects homologs with BLAST,^48^ and matches to CARD’s database and model cut off values.

### ResFinder

ResFinder^32^ v4.0 was available both as a web-based application or the command line. We used ResFinder 4 in this study, which was specifically designed for detecting genotypic resistance in phenotypically resistant samples. ResFinder aligned reads directly to its own curated database without need for assembly.

### deepARG

deepARG^37^ v.2.0 used a supervised deep learning based approach for antibiotic resistance gene annotation of metagenomic sequences. It combines three databases—CARD, ARDB, and UNIPROT—and categorizes them into resistance categories.

### sraX

sraX^36^ v.1.5 was built as a one step tool; in a single command, sraX downloaded a database and aligned contigs to this database with DIAMOND^49^. By default, sraX used CARD, though other options can be specified. As we use default settings for all tools, only CARD was used in this study for sraX. It should be noted that the one step aspect is convenient, but can become lengthy if there are multiple runs and databases need to be downloaded multiple times.

### starAMR

starAMR^35,50^ v.0.7.2 used BLAST+^51^ to compare contigs against a combined database with data from ResFinder, PointFinder, and PlasmidFinder.

### AMR Finder Plus

AMR Finder Plus^34^ v.3.9.3 used BLASTX^48^ translated searches and hierarchical tree of gene families to detect AMR genes. The database was derived from the Pathogen Detection Reference Gene Catalog^52^ and was compiled as part of the National Database of Antibiotic Resistant Organisms (NDARO).

### Performance of software on synthetic metagenomes with high- and -low-prevalence of AMR phenotypes

Each software tool was run against a synthetic mock community of 8 bacteria at three coverage levels that expressed 43 antibiotic resistance phenotypes. Overall, the number of AMR genes detected across all tools ranged from 13 to over 700 at 100x coverage (**Table 3**). For some tools, genes detected did not correspond to a tested phenotype in the mock community, so the prediction fell into the “unknown” category. Among the tools tested, AMR Finder Plus had the highest degree of unclassifiable/unknown results (observed AMR gene not tested in the mock community; **Figure 3**). An overview of these results are available in **Figure 2A**.

**Figure 1:**
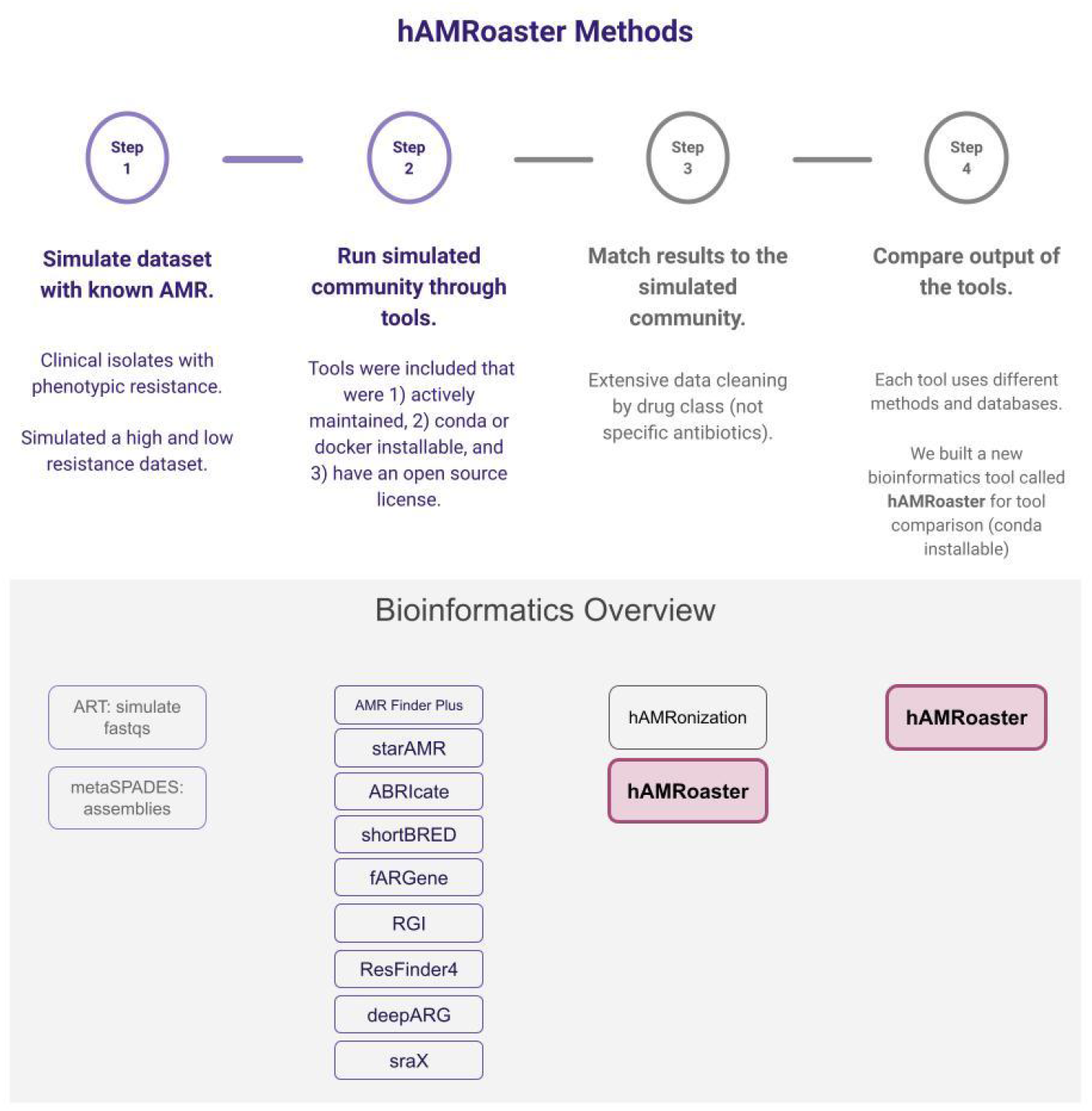
Schematic l Methods.

**Figure 2:**
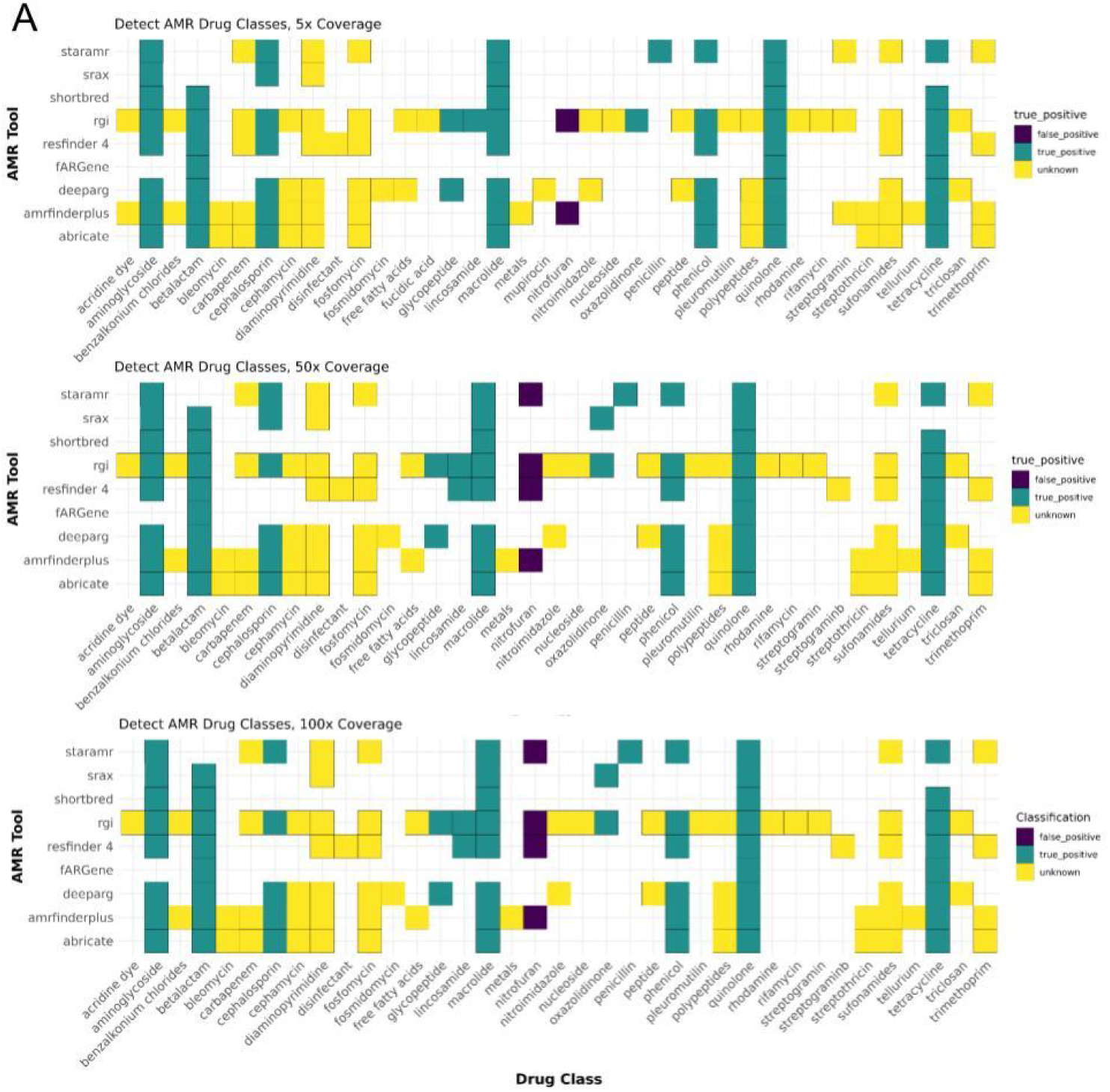
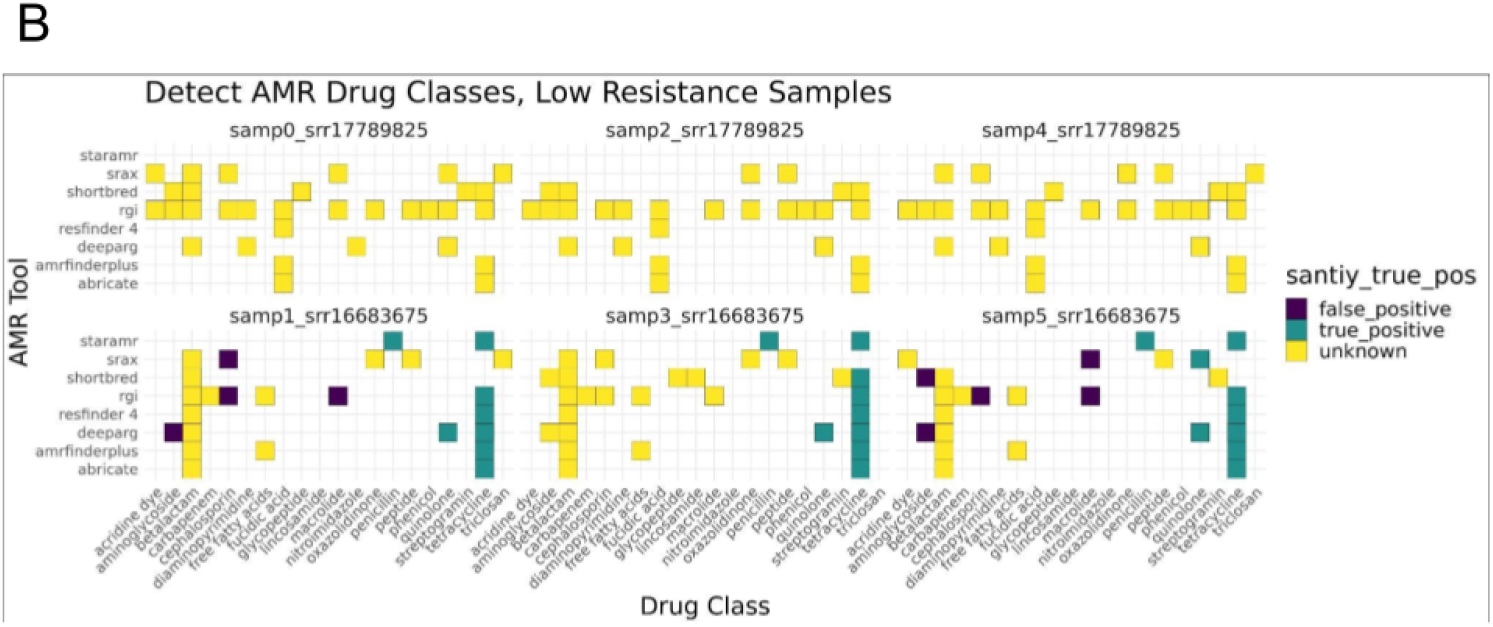
Antimicrobial Resistance (AMR) Genes Detected By Software Tools by Drug Class. AMR Genes detected by each tool across coverage levels, grouped into drug class to which the genes confer resistance with the color coding indicating whether the detection was true positive (green), false positive (purple) or unknown (yellow). Clear spaces in the plot indicate that AMR genes were not detected for the drug class on the x-axis by the tool on the y-axis. Plot A contains the high AMR Data, while plot B contains the low AMR data.

**Figure 3:**
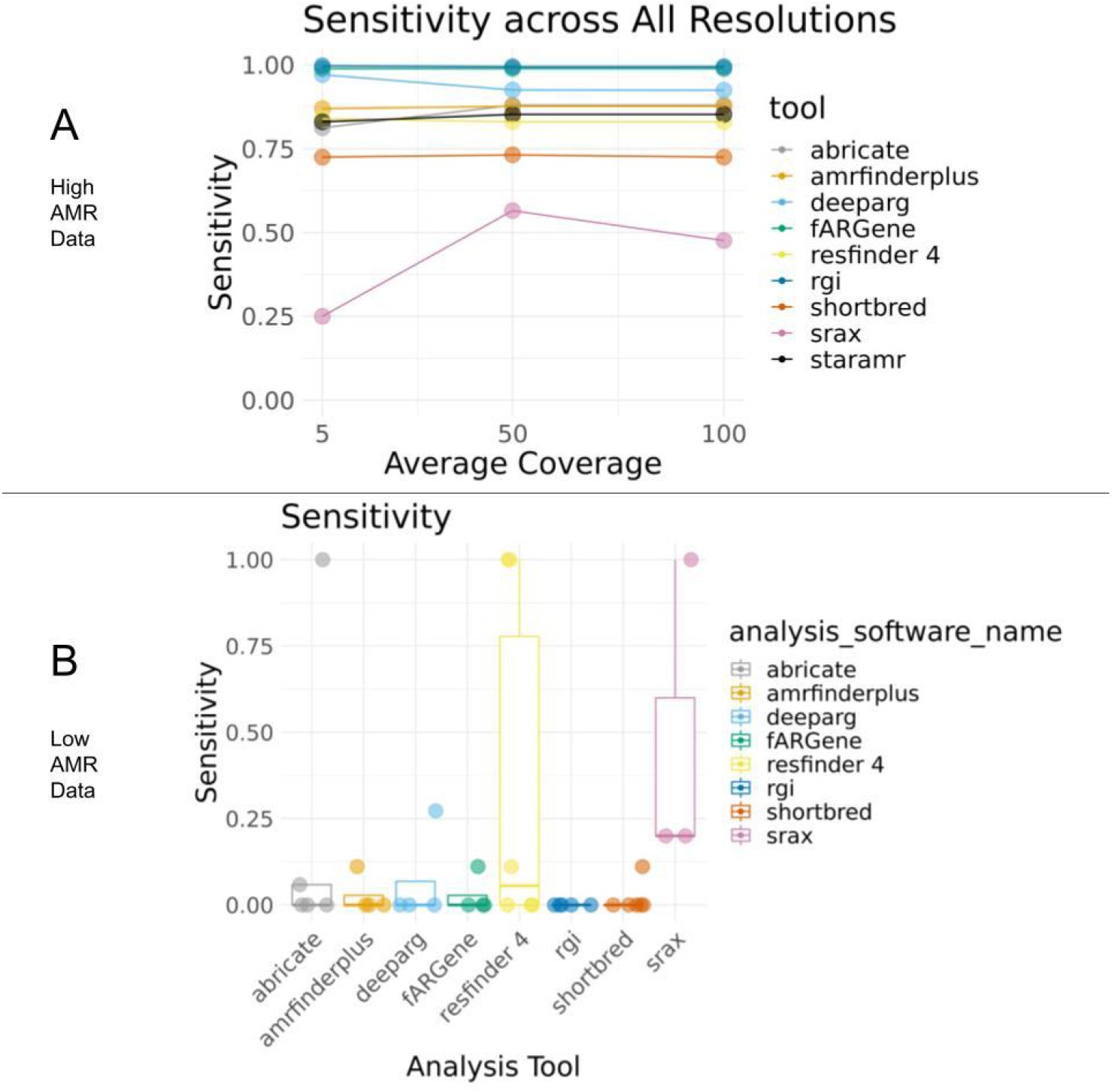
Sensitivity of Software Tools for Detection of Antimicrobial Resistance (AMR) Genes Across Coverage Levels. Sensitivity was calculated as (true positives) / (true positives + false negatives). Most tools were highly sensitive (greater than 0.80). All genes corresponding to “Other” or “Unknown” drug classes were not included in these calculations. Similarly, AMR genes corresponding to phenotypic resistance that was not tested in the mock community was considered “Unknown” and not included in the sensitivity analysis.

After filtering out the AMR genes detected in the simulated human metagenomes (for which AMR phenotypes were unknown), detected AMR genes were examined per sample. None of the tools detected true or false positives for one of the AMR isolates in the low resistance samples (**Figure 2b**). Fewer genes were detected overall compared to the highly resistant sample, as expected for samples with a limited resistance phenotype (**Table 3**), though many of these corresponded to unknown AMR phenotypes and not those included in susceptibility testing.

### Sensitivity and Specificity

Sensitivity tests what portion of AMR genes are correctly identified by a tool when phenotypic resistance to the drug class that gene confers resistance to is present in the mock community. Specificity tests what portion of known negatives (i.e. susceptible drugs from phenotypic testing) do not have AMR genes detected for that drug class. Sensitivity for phenotype detection ranged from >0.99 (RGI) to 0.23 (sraX) at the lowest coverage levels for the highly resistant, antibiotic resistance gene (ARG)-rich dataset sample (**Fig. 2a**). In general, genome coverage did not greatly affect sensitivity, with the exception of sraX, which increased to 0.53 at the highest level. fARGene and deepARG had a high sensitivity value (>0.90) at all coverage levels. RGI, deepARG, and fARGene are all tools that compare reads to a model of AMR instead of aligning reads directly to a database, indicating that this method may be appropriate when high sensitivity values are preferred. As a note, in this ARG-rich dataset, there were only 2 possible true negatives because only two drug classes were always susceptible to antibiotics in those two drug classes when tested (nitrofuran and polypeptide).

In samples with lower numbers of resistance genes, sensitivity and specificity were variable within- and across-tools for samples, with sensitivity much lower than the high resistance community ((0 - <0.45; **Fig. 1b)**Specificity was much higher overall, though variable across samples depending on whether any true positives were detected by the tools (**Table 3**). Precision was highly variable across tools with no consistent trend across tools (range 0 - 1), while accuracy was less variable, with most tools having an accuracy between. 50 and 0.75

### Concordance between tools

An analysis of the agreement between tools of detected resistance to drug classes revealed that overall, agreement was highly variable across tools (0.02 - 0.72 at 5x coverage, **Fig. 5A**) between tools at all coverage levels for the ARG-rich dataset (**Figure 5A, 5B, 5C**). Low agreement was found between most tools in the low AMR samples with the exception of AMR Finder Plus, abricate, and ResFinder4, which had a kappa value > 0.80 (**Figure 5D**).

**Figure 4:**
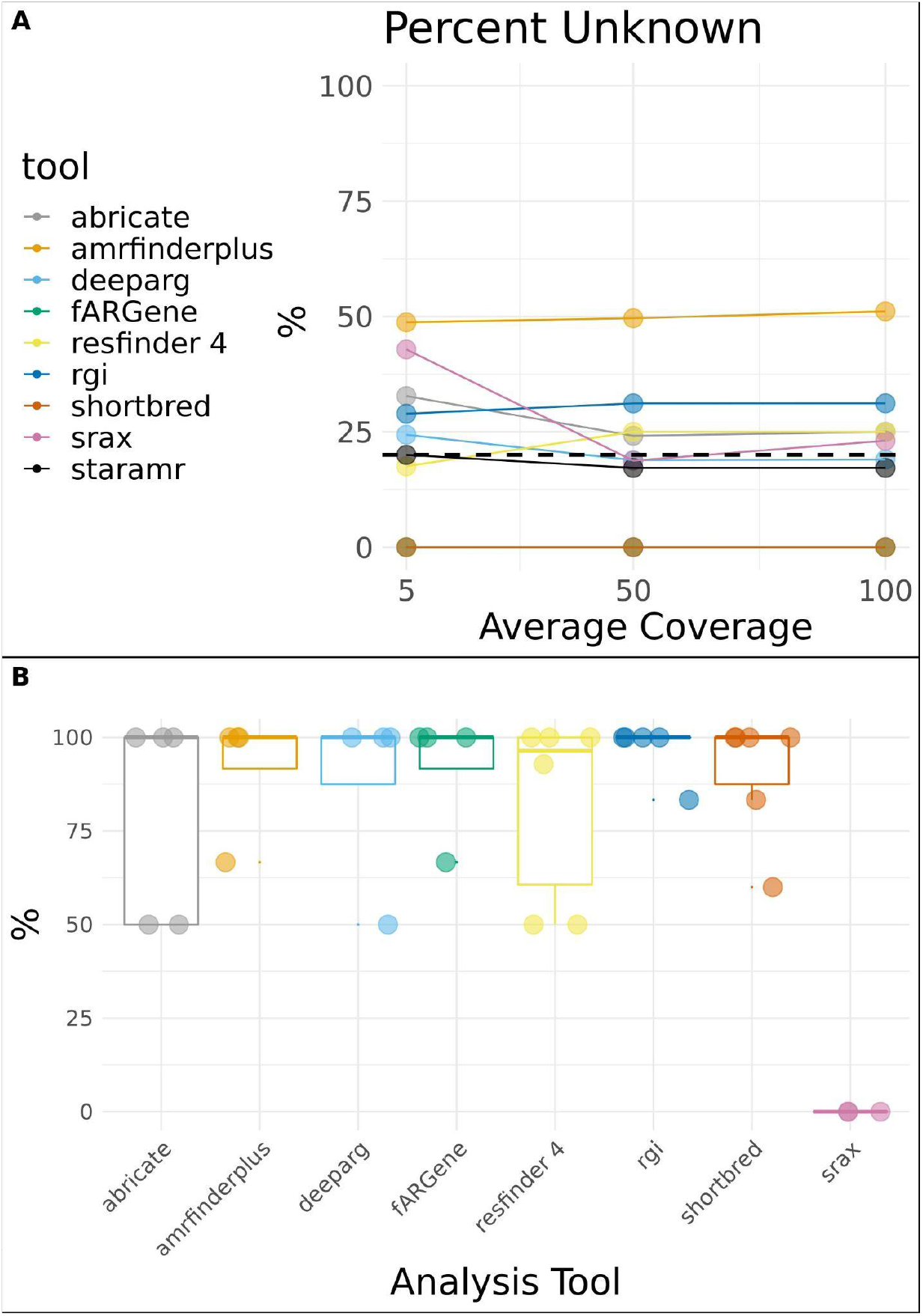
Percent Detection of Unknown Antimicrobial (AMR) Resistance Genes Across Coverage. The percent detection of AMR genes that could not be classified because the drug class the gene confers resistance to was not tested for the high AMR (A) and low AMR (b) data. A black dashed line is placed at 20%, indicating where at least 20% of the detected AMR genes could not be classified.

**Figure 5:**
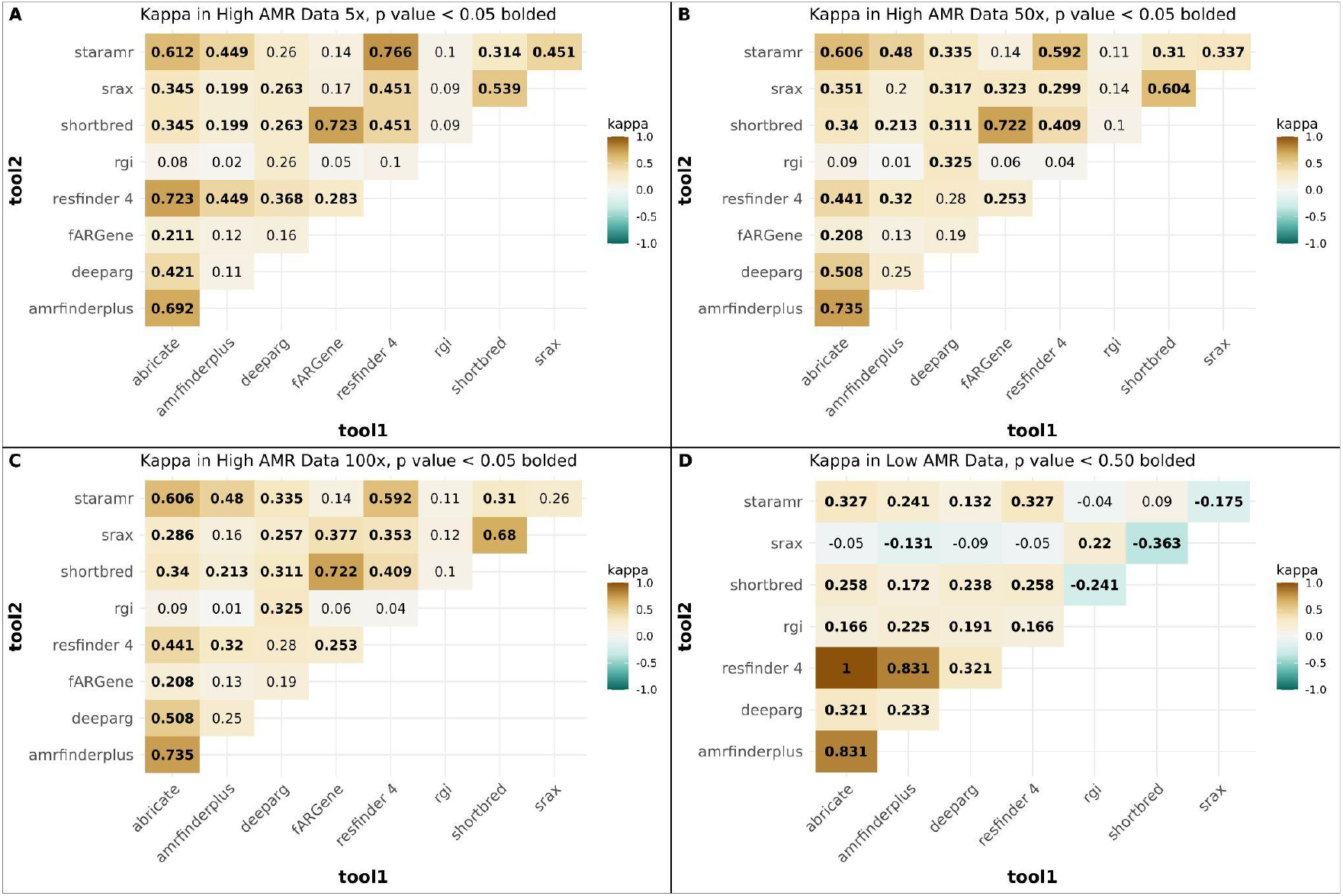
Agreement (Cohen’s Kappa) values between tools across coverage levels calculated in R using the kappa2 function. Agreement between tools in detecting resistance to drug classes is shaded across all plots while kappa values are bolded when the p-value is less than 0.05. A, B, and C display the agreement between tools for the 5x, 50x, and 100x coverage high AMR datasets, respectively. D displays the agreement between tools for the low AMR samples.

## Discussion

### Development of a framework for assessing AMR prediction software performance using synthetic data

There is a considerable research effort to develop new software for predicting AMR using DNA sequence alone. In this dynamic environment, there is a need for researchers and epidemiologists to understand the relative performance of open source software tools. While some tools currently exist for compiling the results of several AMR tools together (hAMRonizer and chARMedDb^53^), this study was motivated by the lack of an open-source pipeline for comparing the results once compiled.

The central challenge in developing this software was to compare detected AMR genes to resistance phenotypes. Detected AMR genes needed to be classified by their corresponding drug class(es) so they could be matched to the known phenotypically resistant drug classes. One hurdle in this translation is that tools use different databases, and some databases classify genes differently. For example, shortBRED classifies gene families, while CARD classifies specific genes. While this analysis checked the drug classification via the DNA/Protein Accession value in CARD, only around 300 of the >1,000 genes detected could directly map to genes in CARD by accession value. The hAMRonization tool overcomes this challenge by providing a drug class column and filling in the values from ChEBI ontology^54^ when possible. The hAMRoaster strategy is to assign a CARD drug class value to every detected AMR gene first by accession number, then by gene name. If neither of these methods assign a drug class for an AMR gene, then the drug class provided by hAMRonization is used. Another challenge in converting detected AMR genes to drug classes is that some drugs are only administered in combination, such as clavulanic acid with amoxicillin. For these instances, resistance to the drug only used in combination (e.g. clavulanic acid) is treated as an “other” drug class and excluded from analysis in hAMRoaster. In these cases, we incorporated the experience of practicing clinicians to identify combination antibiotics into the hAMRoaster antibiotic key.

The analysis presented here used synthetic data to compare tool performance. Synthetic data has the benefit of allowing controlled input with known ground truth. Therefore users can focus on the types of organisms and phenotypes they need to to detect in their own datasets, perform experiments with real samples, and manipulate a range of factors such as relative abundance and sequencing error. The NCBI BioSample repository (used in this study) is an invaluable resource for creating such datasets as it contains many samples with AMR phenotypes determined by international standards. Researchers could also sequence and phenotype culturable organisms in their own laboratories to provide testing standards to evaluate software. Here, we exclusively examined synthetic short read Illumina data, but this analysis strategy could be adapted to understand the effect of using data generated on long read technologies such as the Pacific Bioscience and Oxford Nanopore platforms.

### Overall trends in performance and reasons for variability between tools

We found the sensitivity of almost all tools to be very good in a highly resistant sample (>0.80), with the exception of sraX, which had a proportionally high number of false negatives compared to true positives. However, sensitivity was lower in low-resistance samples (0 - <0.45), indicating that tool selection plays an important role in results for targeted AMR studies. All tools except shortBRED and starAMR detected a large number of genes that were not associated with a lab-determined phenotype in our highly resistant mock community, while this was true for all tools except starAMR in the low-resistance sample. In practice, researchers and epidemiologists may be only interested in a narrow range of AMR phenotypes. Overall, these results indicate when researchers are interested in resistance to a particular drug class as opposed to resistance to a broad range of drug classes, tool selection becomes very important.

We calculated Cohen’s kappa to capture the agreement at the drug class level between AMR tools to see if all AMR tools detected resistance to the same drug classes across samples. We found that agreement at the drug class level was surprisingly low across all tools in the high and low resistance data, though some pairs of tools have higher agreement than others (e.g., AMR Finder Plus, abricate, and ResFinder4 in the low resistance samples; **Figure 5**), indicating that some tools may be better suited for detecting different types of resistance. As such, hAMRoaster provided a table with the number of genes detected per drug class for each tool that may help researchers in selecting an AMR gene detection tool that is best suited for their research question.

This research underlines the need for the further development of software tools for the detection of AMR genes in the human microbiome. It is increasingly recognized that the confined location and genetic diversity of this microbial population provides ideal conditions for genetic exchange among residential microbes and between residential and transient microbes, including pathogenic microbes. Notably, rates of horizontal gene transfer among bacteria in the human microbiome (especially the gastrointestinal tract) are estimated to be many times higher than among bacteria in other diverse ecosystems, such as soil.^55^ Refined tools appropriate for use in shotgun metagenomic data will be important for tracking the spread of AMR genes from diverse environmental sources to the human microbiome and across sites in the human body and understanding whether AMR genes are derived from vertical inheritance or via horizontal gene transfer.

In conclusion, this study compared bioinformatics tools for detecting AMR genes in a simulated short read metagenomic sample at three coverage levels at one time point. While tools use slightly different methods and databases, these tools overall had high sensitivity for detection of AMR genes. Moreover, agreement between tools was sometimes low, indicating the importance of careful tool selection. We advocate that researchers should test these software tools using pipelines such as hAMRoaster with a synthetic community that highlights the resistance profiles and sample of interest.

## Acknowledgements

We thank Jon Moller for helping to create the hAMRoaster name.

---

Supplementary text 1: URL link to tweet https://twitter.com/emily_wissel/status/1336013892116488195

Supplementary table 1: tidy table of data https://does.google.com/spreadsheets/d/1bfACqEh0nkS65vCUi5DfMg4PvW0fHxbtrv0PgKtlgT4/edit#gid=53644837

